# Robust Genome Editing of Single-Base PAM Targets with Engineered ScCas9 Variants

**DOI:** 10.1101/620351

**Authors:** Pranam Chatterjee, Noah Jakimo, Joseph M. Jacobson

## Abstract

Programmable CRISPR enzymes are powerful and versatile tools for genome editing. They, however, require a specific protospacer adjacent motif (PAM) flanking the target site, which constrains the accessible sequence space for position-specific genome editing applications, such as base editing and homology-directed repair. For example, the standard Cas9 from *Streptococcus pyogenes* requires a PAM sequence of 5’-NGG-3’ downstream of its RNA-programmed target. Recently, three separate Cas9 enzymes (xCas9-3.7, SpCas9-NG, and ScCas9) have been independently engineered or discovered to reduce the PAM specificity to a single guanine (G) nucleotide, thus greatly expanding the number of targetable sequences. In this study, we have employed motifs from closely-related orthologs to engineer and optimize ScCas9 to exhibit enhanced genome editing and higher fidelity. Our engineered variants demonstrate superior activity within gene repression and nucleolytic contexts and possess effective base editing capabilities.

## Introduction

RNA-guided endonucleases of the CRISPR-Cas bacterial immune system have been successfully harnessed for various genome editing and regulation applications (*1–3*). To access specific targets, however, these enzymes require a protospacer adjacent motif (PAM), which is determined by DNA-protein interactions, to immediately follow the DNA sequence specified by the single guide RNA (sgRNA) (*1, 4–7*). The Cas9 from *Streptococcus pyogenes* (SpCas9), for example, requires an 5’-NGG-3’ motif downstream of its RNA-programmed DNA target (*1, 6, 7*), severely restricting position-specific genome editing applications, such as base editing (*8, 9*) and homology-directed repair (*10*), which represent promising routes for effective therapeutics and biotechnologies.

To relax this constraint, additional Cas variants with distinct PAM requirements have been either discovered (*11–14*) or engineered (*15–18*). In fact, within the past year, three groups have independently reduced the 5’-NGG-3’ PAM specificity of SpCas9 to a single guanine (G) nucleotide, by employing phage-assisted continuous evolution (xCas9-3.7) (*19*), structure-guided rational design (SpCas9-NG) (*20*), and bioinformatics-mediated ortholog discovery (ScCas9) (*21*). Together, these enzymes increase the targetable DNA sequence space to nearly 50%.

While these three new tools represent an exciting expansion of targets for genome editing, they each present their own shortcomings and limitations. For example, SpCas9-NG demonstrates reduced efficiency on 5’-NGC-3’ PAM targets (*20*), while ScCas9 is notably inefficient at modifying target sequences within different gene contexts (*21*). Finally, xCas9-3.7 has been suggested to possess higher fidelity rather than broad PAM recognition (*22, 23*). Thus, there is a critical need for continual improvement of these enzymes for genome editing purposes.

In this study, we engineer broad-targeting and efficient ScCas9 enzymes (Sc+ and Sc++), by utilizing evolutionary information from closely-related orthologs to generate two novel modifications to the original ORF. Taken together, these alterations enable Sc+ and Sc++ to possess enhanced editing capabilities in both bacterial and human cells, in comparison to SpCas9, xCas9-3.7, SpCas9-NG, and ScCas9. Finally, we engineer a high-fidelity variant of Sc++ for genome modification and demonstrate its improved specificity.

## Results

### Engineering of ScCas9 Variants

SpCas9-NG and xCas9-3.7 both harbor various substitutions in their open reading frames (ORFs) that allow reduced specificity from the canonical 5’-NGG-3’ to the more minimal 5’-NGN-3’ PAM. Specifically, positions 1218-1219 for both enzymes have been shown to be the most consequential in terms of PAM recognition (*20, 24*). To engineer ScCas9 to possess improved PAM targeting capabilities, we performed global pairwise alignments using the BLOSUM62 scoring matrix (*25*) of various *Streptococcus* Cas9 orthologs to SpCas9, xCas9-3.7, and SpCas9-NG at these critical residues. Our sequence alignment isolated a positive-charged lysine residue, derived from the *S. gordonii* Cas9 ORF. Substituting positive-charged residues into the PAM-interacting domain (PID) of Cas enzymes has been suggested to allow for the formation of novel PAM-proximal DNA contacts (*18*). Motivated by this finding, we thus substituted the corresponding T1227K mutation into the ORF of ScCas9, generating ScCas9+ (Sc+) (Figure 1A).

**Figure 1:**
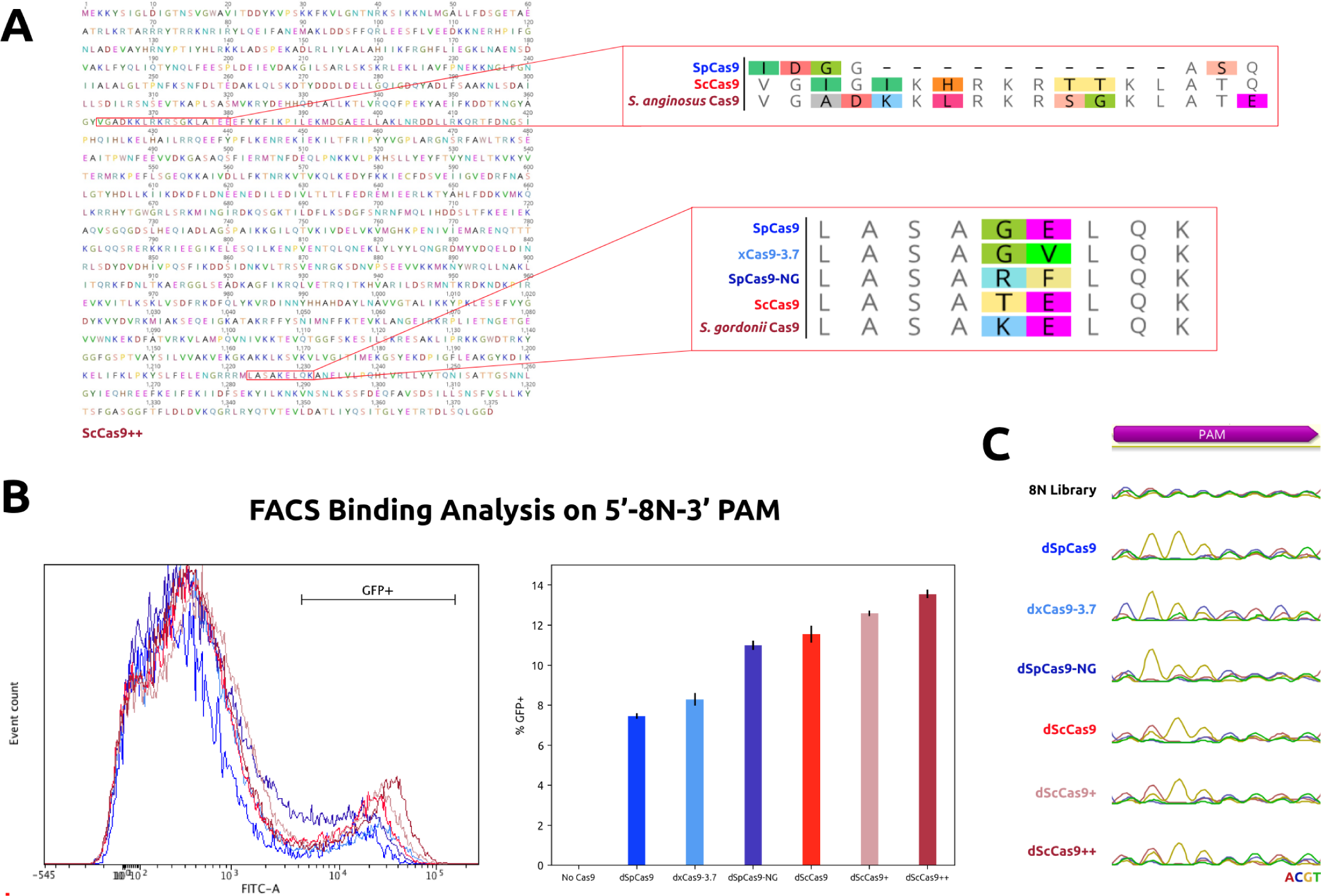
Engineering and PAM Determination of Improved ScCas9 Variants. A) Amino acid sequence of Sc++. SpCas9, SpCas9-NG, xCas9-3.7, and ScCas9 were aligned with various *Streptococcus* Cas9 orthologs, employing the BLOSUM62 scoring matrix, to identify the T1227K mutation derived from *Streptococcus gordonii*. Sequence alignment of ScCas9 with various *Streptococcus* Cas9 orthologs further isolated a novel loop structure from *Streptococcus anginosus* harboring an additional lysine residue and a flexible “SG” motif. B) PAM binding analysis of single G PAM Cas9 variants on a 5’-NNNNNNNN-3’ (8N) PAM library. Each dCas9 plasmid was electroporated in duplicates, subjected to FACS analysis, and gated for GFP expression. Subsequently, percentages of GFP-positive cells were averaged. Standard deviation was used to calculate error bars. C) PAM binding enrichment visualization. PAM profiles are represented by DNA chromatograms via amplification of PAM region following plasmid extraction of GFP-positive *E. coli* cells and subsequent Sanger sequencing.

One of the defining characteristics of ScCas9’s PAM flexibility is its employment of a positive-charged loop, in positions 367 to 376 of its ORF, which does not exist in SpCas9 or its engineered variants (*21*). Our sequence alignments identified a divergent insertion from *S. anginosus*, which not only maintains the positive charge of the ScCas9 loop by compensating an extra lysine residue for a histidine, but also possesses an “SG” motif, a flexible sequence of residues used for linker design in protein engineering (*26*) (Figure 1A). We therefore hypothesized that this novel loop may improve the targeting capabilities and efficiency of ScCas9 by allowing for more flexible protein-phosphate backbone contacts with the PAM sequence. Thus, we substituted the loop sequence from *S. anginosus* into the Sc+ ORF to generate ScCas9++ (Sc++) (Figure 1A).

### Determination of PAM Sequences Recognized by Engineered Sc-Cas9 Variants

To comprehensively profile the PAM specificity of Sc+ and Sc++, in comparison to SpCas9, xCas9-3.7, and SpCas9-NG, as well as the wild-type ScCas9, we utilized a previously-developed positive selection bacterial screen based on green fluorescent protein (GFP) expression conditioned on PAM binding, termed PAM-SCANR (*27*). Following transformation of the PAM-SCANR plasmid, harboring a randomized 5’-NNNNNNNN-3’ (8N) PAM library, an sgRNA plasmid targeting the fixed PAM-SCANR protospacer, and a corresponding dCas9 plasmid, we conducted FACS analysis to first determine the percent of GFP-positive cells in each population, a relative proxy for the percent of total PAM sequences being bound. Our results demonstrated that both dSc+ and dSc++ bind to a greater percentage of PAM sequences, and dSc++ exhibits a shifted GFP-positive population, suggesting stronger binding capabilities and improved efficiency (Figure 1B). Plasmid DNA from FACS-sorted GFP-positive cells and presorted cells were then extracted and amplified, and enriched PAM sequences were identified by Sanger sequencing, and visualized utilizing DNA chromatograms. Sequencing results indicate that the ScCas9 variants possess improved PAM specificity, as compared to xCas9-3.7, which demonstrates notable dependence on bases in downstream positions, and SpCas9-NG, which may require additional G nucleotides in positions 3 or 4 for efficient binding (Figure 1C). While exhibiting similar specificity to ScCas9 and Sc+, Sc++ comparatively enjoys greater independence at position 4 in the PAM sequence (Figure 1C). Taken together, these results suggest that Sc+ and Sc++ possess broader targeting capabilities and, potentially, enhanced efficiency for genome editing applications, thus prompting their characterization in human cells.

### Genome Editing Capability of Engineered ScCas9 Variants

We compared the PAM specificities and nucleolytic capabilities of Sc+ and Sc++ to SpCas9, xCas9-3.7, SpCas9-NG, and ScCas9 by transfecting HEK293T cells with plasmids expressing each variant individually alongside one of 16 sgRNAs, together directed to four genomic loci with diverse PAM sequences, collectively representing every base at each position in the PAM window (Table 1). The sgRNA sequences were shifted by one base for xCas9-3.7 and SpCas9-NG to account for their reported 5’-NGN-3’ PAM preferences, so to equivalently compare these enzymes to ScCas9 variants with 5’-NNG-3’ specificities. After 5 days post-transfection, indel formation was quantified from Sanger sequencing ab1 files using the TIDE algorithm (*28*) following PCR amplification of the target genomic region. Our results demonstrate that Sc+ and Sc++ can effectively edit across the various genomic loci, and demonstrate improved indel formation percentages for a majority of the targets tested (Figure 2A). SpCas9, xCas9-3.7, and SpCas9-NG all edit on “GG” PAM targets, and maintain activity on various 5’-AGN-3’ PAM sequences. xCas9-3.7 and SpCas9-NG additionally edit few sites that harbor 5’-CGN-3’ and 5’-TGN-3’ sequences, but perform poorly on all tested 5’-NGC-3’ PAM targets, consistent with previously reported data (*19, 20, 22–24*). Sc+ and Sc++, on the other hand, improve greatly upon the editing capabilities of the wild-type ScCas9 enzyme, demonstrating nearly 3-fold improvement in indel formation efficiency on certain 5’-NNGC-3’ targets, and even editing sites at which ScCas9, xCas9-3.7, and SpCas9-NG have negligible activity (Figure 2A). We subsequently incorporated the D10A nickase version of ScCas9+ into the BE3 base editing architecture to examine whether our engineered ScCas9 variants may enable successful C*→*T base conversion. Following transfection of the ScCas9+ BE3 plasmid and plasmids encoding sgRNAs directed at 4 genomic sites with PAM sequences representing each base at both flanking positions (Table 1), we observed evident C*→*T base editing activities in the 5-nucleotide editing window, in comparison to the unedited control (Figure 2B), demonstrating that our engineered variants can be further utilized for base editing purposes. Together, this data suggests that Sc+ and Sc++ are efficient, broad-targeting enzymes that can be harnessed for diverse genome editing applications.

**Table 1:**
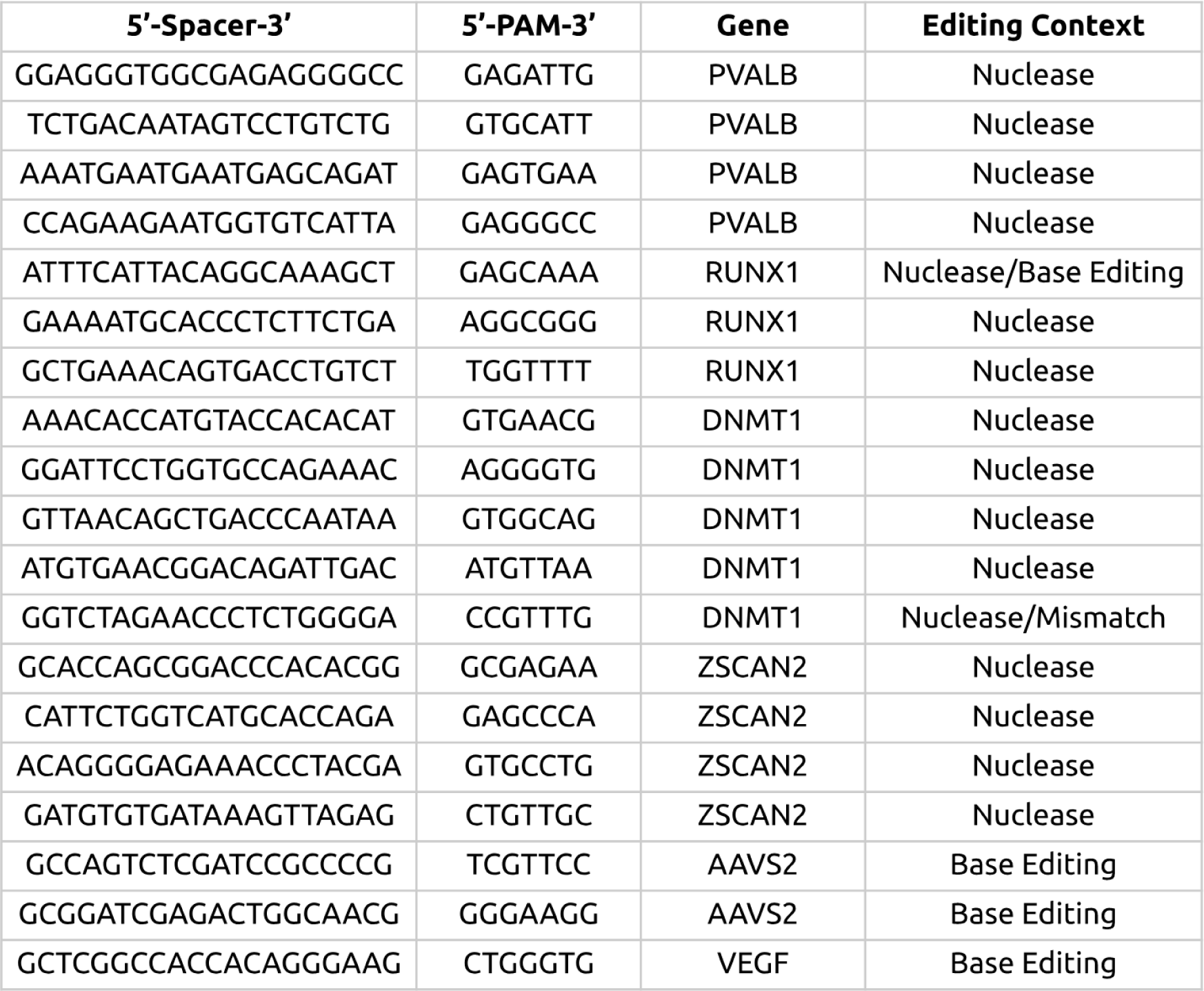
Sequence Information for Genome Editing in Human Cells. Spacer and PAM sequences indicated are for use with ScCas9 variants and the standard SpCas9. All sequences for xCas9-3.7 and SpCas9-NG are shifted one base in the 3’ direction for equivalent comparison purposes, due to their reported 5’-NGN-3’ PAM sequences.

**Figure 2:**
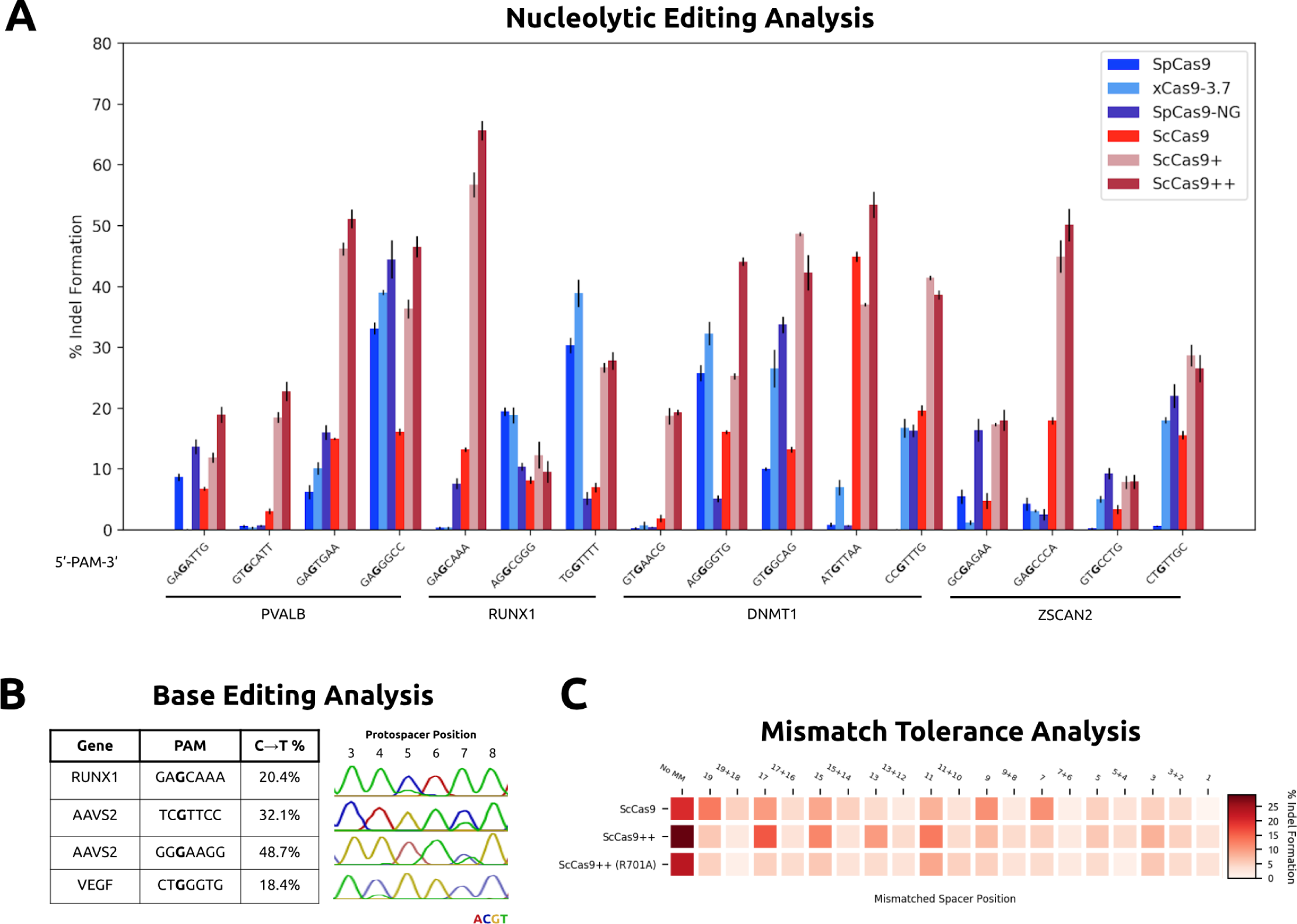
Genome Editing Capabilities of Engineered ScCas9 Variants. A) Quantitative analysis of nucleolytic editing with single G PAM Cas9 variants. Indel frequencies were determined via the TIDE algorithm following PCR amplification of indicated genomic loci, in comparison to unedited controls for each gene target. All samples were performed in duplicates and quantified indel formation values were averaged. Standard deviation was used to calculate error bars. B) Quantitative analysis of C *→* T base editing with ScCas9+ BE3. C *→* T conversion frequencies were determined via the BEEP algorithm, in comparison to unedited controls, following PCR amplification of targeted genomic loci. All samples were performed in duplicates and quantified base editing values were averaged. C) Efficiency heatmap of mismatch tolerance assay on a genomic target. Quantified indel frequencies, as assessed by the TIDE algorithm, are exhibited for each labeled single or double mismatch in the sgRNA sequence for the indicated Cas9 variant. The target protospacer sequence within the DNMT1 gene is 5’-GGTCTAGAACCCTCTGGGGA-3’, possessing a PAM sequence of 5’-CCGTTTG-3’ (Table 1).

### Mismatch Tolerance Profile of a High-Fidelity Sc++ Nuclease

To assess the off-target propensity of our engineered nucleases, we conducted a mismatch tolerance assay (*29*) employing sgRNAs harboring double or single mismatches to a fixed protospacer in the endogenous DNMT1 gene with a non-canonical 5’-CCGT-3’ PAM sequence (Table 1). Following TIDE analysis, we observed that ScCas9 and Sc++ share similar mismatch tolerance profiles across the spacer sequence (Figure 2C). Overall, double mismatches are tolerated less than single mismatches, and mismatches within the PAM-distal region of the spacer generally allow higher editing rates. As Sc++ possesses higher efficiency overall, however, the magnitude of activity for mismatched spacer sequences is greater (Figure 2C). Thus, to ameliorate the mismatch tolerance of Sc++, we engineered a high-fidelity variant harboring the R701A mutation, which was previously isolated via high-throughput bacterial selection for SpCas9 to maintain high on-target activity while reducing off-target editing (*30*). Our engineered variant demonstrated a slight reduction in on-target editing from that of Sc++, but exhibited reduced activity on mismatched sequences (Figure 2C). Overall, these results motivate the usage of this high-fidelity Sc++ for broad and efficient genome editing with reduced mismatch tolerance.

## Discussion

The application of CRISPR-Cas9 has been hampered by the inaccessibility of genomic sequences, largely due to the PAM restriction. The recent discoveries of ScCas9, xCas9-3.7, and SpCas9-NG, all reporting to possess single G PAM specificity, significantly increased the targetable space, potentially allowing for expanded base editing activities, more efficient homology-directed repair, and denser screening platforms. As all have been shown to possess limitations, however, including inefficient targeting of certain single G PAM sequences, we have engineered ScCas9 to possess increased efficiency and broader targeting capabilities, by utilizing sequence information from engineered Cas9 variants as well as uncharacterized *Streptococcus* Cas9 orthologs. Sc+ and Sc++ nucleases outperform SpCas9, xCas9-3.7, SpCas9-NG, and ScCas9 as genome editing tools, and can thus be harnessed for various applications, including base editing. Furthermore, we have demonstrated that, due to high sequence homology of ScCas9 and SpCas9, previous modifications made to SpCas9, such as high-fidelity mutations (*30*), can be ported into these engineered variants for improved functionality. Nonetheless, it remains critical to continue improving these enzymes to enable versatile genome engineering and eventually provide access to every sequence in the entire genome. Sc+ and Sc++, with their broad targeting range and high genome editing efficiency, will hopefully serve as platforms toward this goal.

## Materials and Methods

### Generation of Plasmids

The pX330-SpCas9-NG (Addgene Plasmid #117919) and xCas9 3.7 (Addgene Plasmid #108379) were gifts from Osamu Nureki and David Liu, respectively. The Cas9 from *S. canis* was codon optimized for human cell expression, ordered as multiple gBlocks from Integrated DNA Technologies (IDT), and assembled using Gibson Assembly into a mammalian expression backbone harboring an EF1*α* promoter and coexpressing GFP. Engineering of the coding sequence of ScCas9 to generate the T1227K, *S. anginosus* loop, and R701A substitutions were conducted using the KLD Enzyme Mix (NEB) following PCR amplification with mutagenic primers (Genewiz). To assemble ScCas9 base editing plasmids, pCMV-BE3 (Addgene Plasmid #73021) was received as a gift from David Liu. Similarly, the ORF of the ScCas9+ D10A nickase was amplified by PCR to generate 35 bp extensions for subsequent Gibson Assembly into each base editing architecture backbone. sgRNA plasmids were constructed by annealing oligonucleotides coding for crRNA sequences (Table S1) as well as 4 bp overhangs, and subsequently performing a T4 DNA Ligase-mediated ligation reaction into a plasmid backbone immediately downstream of the human U6 promoter sequence. Assembled constructs were transformed into 50 *µ*L NEB Turbo Competent *E. coli* cells, and plated onto LB agar supplemented with the appropriate antibiotic for subsequent sequence verification of colonies and plasmid purification.

### PAM-SCANR Assay

Plasmids for the SpCas9 sgRNA and PAM-SCANR genetic circuit, as well as BW25113 ΔlacI cells, were generously provided by the Beisel Lab (North Carolina State University). Plasmid libraries containing the target sequence followed by either a fully-randomized 8-bp 5’-NNNNNNNN-3’ library or fixed PAM sequences were constructed by conducting site-directed mutagenesis, utilizing the KLD enzyme mix (NEB) after plasmid amplification, on the PAM-SCANR plasmid flanking the protospacer sequence (5’- CGAAAGGTTTTGCACTCGAC-3’). Nuclease-deficient mutations (D10A and H850A) were introduced to the ScCas9 variants using Gibson Assembly as previously described. The provided BW25113 cells were made electrocompetent using standard glycerol wash and resuspension protocols. The PAM library and sgRNA plasmids, with resistance to kanamycin (Kan) and carbenicillin (Crb) respectively, were co-electroporated into the electrocompetent cells at 2.4 kV, outgrown, and recovered in Kan+Crb Luria Broth (LB) media overnight. The outgrowth was diluted 1:100, grown to ABS600 of 0.6 in Kan+Crb LB liquid media, and made electrocompetent. Indicated dCas9 plasmids, with resistance to chloramphenicol (Chl), were electroporated in duplicates into the electrocompetent cells harboring both the PAM library and sgRNA plasmids, outgrown, and collected in 5 mL Kan+Crb+Chl LB media. Overnight cultures were diluted to an ABS600 of 0.01 and cultured to an OD600 of 0.2. Cultures were analyzed and sorted on a FACSAria machine (Becton Dickinson). Events were gated based on forward scatter and side scatter and fluorescence was measured in the FITC channel (488 nm laser for excitation, 530/30 filter for detection), with at least 10,000 gated events for data analysis. Sorted GFP-positive cells were grown to sufficient density, and plasmids from the pre-sorted and sorted populations were then isolated, and the region flanking the nucleotide library was PCR amplified and submitted for Sanger sequencing (Genewiz). Additional details on the PAM-SCANR assay can be found in Leenay, et al. (*27*).

### Cell Culture and DNA Modification Analysis

HEK293T cells were maintained in DMEM supplemented with 100 units/ml penicillin, 100 mg/ml streptomycin, and 10% fetal bovine serum (FBS). sgRNA plasmids (100 ng) and effector (nuclease and BE3) plasmids (100 ng) were transfected into cells as duplicates (2 × 10^4^/well in a 96-well plate) with Lipofectamine 3000 (Invitrogen) in Opti-MEM (Gibco). After 5 days post-transfection, genomic DNA was extracted using QuickExtract Solution (Epicentre), and genomic loci were amplified by PCR utilizing the Phusion Hot Start Flex DNA Polymerase (NEB). Amplicons were enzymatically purified and submitted for Sanger sequencing (Genewiz). Sanger sequencing ab1 files were either analyzed using the TIDE algorithm (tide.deskgen.com) in comparison to an unedited control to calculate indel frequencies, or by the internally-developed BEEP software (github.com/mitmedialab/BEEP) for base editing analysis. All samples were performed in duplicates and modification values were averaged. Standard deviation was used to calculate error bars.

## Acknowledgments

We thank Dr. Ed Boyden for access to cell culture, in addition to Dr. Neil Gershenfeld and Dr. Shuguang Zhang for shared lab equipment. We further thank Megi Topalli and Andrew Hennes for technical assistance.

## Author Contributions

P.C. and N.J. conceived study and designed experiments. P.C. carried out experiments and conducted data analyses. P.C. wrote the paper. P.C, N.J., and J.M.J. reviewed the paper. J.M.J. supervised the project.

## Declarations

This work was supported by the consortia of sponsors of the MIT Media Lab and the MIT Center for Bits and Atoms. The authors have filed a patent application related to this work.

## Data and Materials Availability

All data needed to evaluate the conclusions in the paper are present in the paper. Additional data and materials related to this paper may be obtained from the authors upon reasonable request. The pScMAX+ plasmid, a codon optimized mammalian expression vector encoding the ScCas9+ nuclease, is currently available on Addgene (Plasmid #124698).

